# Molecular docking of Pyocyanin from *Pseudomonas aeruginosa* with NADPH oxidase and Toll-like receptor 4 : Exploring the Link Between Oral Microbiome and Oral Cancer Pathogenesis

**DOI:** 10.1101/2025.08.24.671957

**Authors:** Sheilina Choudhary, Prince Kumar, Suman Sundar Mohanty, Praveen Kumar Anand, Janesh Kumar Gautam

**Author notes:** Equal First Author (Sheilina Choudhary, Prince Kumar).

## Abstract

Oral cancer is among the leading cancers in India, with its progression influenced not only by genetic and environmental factors but also by dysbiosis in the oral microbiome. *Pseudomonas aeruginosa*, an opportunistic Gram-negative bacterium, has been implicated in oral carcinogenesis through the action of its virulence factor, pyocyanin—a redox-active phenazine that induces reactive oxygen species (ROS), promoting oxidative stress, immune dysregulation, and inflammation. Despite evidence linking pyocyanin to ROS-related pathways, its molecular interaction with key host proteins like NADPH oxidase and Toll-like receptor 4 (TLR4) remains unexplored.

The present study aims to investigate the molecular interactions of pyocyanin with NADPH oxidase and TLR4 through molecular docking, aiming to elucidate its role in oxidative and inflammatory mechanisms associated with oral cancer pathogenesis.

The structures of pyocyanin and the NADPH oxidase (NOX2) and TLR4 target proteins were retrieved from PubChem and RCSB PDB, respectively. CASTp was used to identify potential binding pockets. Molecular docking was performed using the HADDOCK server. The docked complexes were analyzed based on HADDOCK scores, binding energies, RMSD, and buried surface area. Interaction profiles were visualized using BIOVIA Discovery Studio.

Among the docking clusters, Cluster 6 for the NOX2–pyocyanin complex showed a favorable HADDOCK score (84.4 ± 2.9), low RMSD (0.2 ± 0.0), and substantial buried surface area (456.1 ± 13.2 Å^2^). For the TLR4–pyocyanin complex, Cluster 1 exhibited the most favorable interaction profile with a HADDOCK score of 25.0 ± 5.5 and a Z-score of –2.3. Binding site prediction and hydropathy analysis supported the structural feasibility of pyocyanin interaction with both targets.

This study provides structural insights into the potential interaction of pyocyanin with NADPH oxidase and TLR4, supporting its role in modulating oxidative stress and inflammatory signaling pathways. These findings enhance our understanding of oral microbiome–host interactions and their contribution to oral cancer pathogenesis.

## Introduction

Oral cancer ranks as the 16th most common cancer globally. In India, oral cancer ranks as the second most prevalent cancer, with an incidence rate of 10.4% and a mortality rate of 9.3% (Kijowska et al., 2024). The development and progression of oral cancer are influenced by genetic mutations, including alterations in *TP53, CDKN2A (p16), EGFR, NOTCH1*, and *PIK3CA* (Ali et al., 2017). The major risk factors for oral cancer are tobacco and alcohol consumption, Human papillomavirus (HPV) infection, along with poor oral hygiene, prolonged sun exposure, nutritional deficiencies, genetic predisposition, and chronic mechanical irritation (Gautam et al., 2025; Gautam et al., 2025; Smith et al., 2010).

Emerging evidence suggests that the oral microbiome plays a critical role in oral carcinogenesis. Dysbiosis in the oral microbiota may lead to the production of cancer-associated metabolites, fostering a microenvironment conducive to tumor growth (Irfan et al., 2020; Wang et al., 2024; Zhang et al., 2020). Pathogenic bacteria such as *Fusobacterium nucleatum* and *Porphyromonas gingivalis* have been implicated in promoting chronic inflammation, immunosuppression, and tumor invasion (Lekshmi Priya et al., 2025). These bacteria can activate immune receptors, particularly Toll-like receptors (TLRs), thereby modulating host immune responses and facilitating tumor progression (Tagg et al., 2025). In contrast, commensal bacteria such as *Streptococcus salivarius* and *Lactobacillus* spp. support mucosal immunity and exhibit antitumor properties (Asoudeh-Fard et al., 2017).

Among pathogenic microbes, *Pseudomonas aeruginosa*, a Gram-negative opportunistic bacterium, has been associated with oral cancer due to its virulence factor pyocyanin (Qin et al., 2022). Pyocyanin is a redox-active phenazine that generates reactive oxygen species (ROS) such as hydrogen peroxide (H_2_O_2_) and superoxide anion (O_2_^−^) by cycling between its oxidized (Pyo^°^) and reduced (Pyo^−^) forms (Reszka et al. 2004). ROS overproduction disrupts key cellular processes including mitochondrial respiration, ER function, and peroxisomal metabolism. Persistent oxidative stress damages DNA, proteins, and lipids, ultimately leading to genomic instability, protein degradation, and lipid peroxidation (Kudryavtseva et al., 2016). Furthermore, ROS accumulation impairs immune function, promoting immune evasion by *P. aeruginosa* (Xin et al., 2023).

One of the key enzymatic sources of ROS is NADPH oxidase, a multi-subunit complex that transfers electrons from NADPH to molecular oxygen. While NADPH oxidase activity is essential for pathogen defense, its dysregulation results in excessive ROS generation, leading to oxidative tissue damage (Tarafdar & Pula, 2018). Pyocyanin has been shown to modulate NADPH oxidase by accepting electrons from NADPH, thereby amplifying ROS production (Rada et al., 2013). This further promotes oxidative stress-mediated cellular injury and inflammatory signaling.

ROS also serve as upstream activators of innate immune receptors such as Toll-like receptor 4 (TLR4), which recognizes lipopolysaccharides (LPS) from Gram-negative bacteria. TLR4 signaling occurs via two pathways: the TRIF-dependent pathway, which induces type I interferons via IRF3, and the MyD88-dependent pathway, which activates NF-κB and promotes the release of pro-inflammatory cytokines (Ullah et al., 2016). ROS-dependent activation of TLR4 establishes a proinflammatory feedback loop, contributing to chronic inflammation, immune dysregulation, and tumor promotion (González-Carnicero et al., 2023; Kong et al., 2010; Sabroe et al., 2005).

Previous studies have suggested that pyocyanin regulates NADPH oxidase activity and induces ROS generation (Rada et al., 2013), and that NADPH oxidase–dependent ROS can modulate TLR4 signaling, contributing to oral carcinogenesis (Park & Lee, 2013). However, the direct molecular interaction of pyocyanin with NADPH oxidase and TLR4 has not been clearly elucidated. In this study, we investigate the potential interactions between pyocyanin and two key host proteins—NADPH oxidase and TLR4—using in silico molecular docking approaches. Our findings aim to elucidate the structural basis of pyocyanin-mediated modulation of oxidative and inflammatory pathways involved in oral cancer pathogenesis.

## Methods

### 1. Ligand Preparation

The three-dimensional (3D) structure of Pyocyanin was retrieved from the PubChem database [PubChem CID: 6817]. The structure was downloaded in SDF format and subsequently converted to PDB format using BIOVIA Discovery Studio. The molecule was energy-minimized using the BIOVIA Discovery Studio to obtain an optimized conformation suitable for molecular docking.

### 2. Target Preparation

The 3D structures of NADPH oxidase (NOX) [PDB ID-7U8G] and Toll-like receptor 4 (TLR4) [PDB ID-3FXI] were retrieved from the RCSB Protein Data Bank. The NADPH oxidase structure was resolved by electron microscopy at 3.2 Å resolution, while the TLR4 structure was determined by X-ray diffraction at 3.1 Å resolution. The protein structures were downloaded in PDB format.

Protein preparation was carried out using BIOVIA Discovery Studio. Water molecules were removed, and polar hydrogen atoms were added to stabilize the protein structure. The processed protein structures were saved in PDB format and used for molecular docking analyses.

### 3. Identification of Binding Pockets of target proteins

Potential binding sites in the selected protein structures were identified using the Computed Atlas of Surface Topography of Proteins (CASTp) server (http://sts.bioe.uic.edu/castp/index.html). CASTp calculates the volume, area, and geometry of protein surface pockets and internal cavities, enabling precise identification of likely ligand-binding sites.

The most appropriate binding pockets were selected based on parameters such as size, accessibility, and shape complementarity. This ensured that docking was conducted at functionally relevant and structurally favorable sites, increasing the specificity and stability of ligand-protein interaction.

### 4. Hydropathy plot generation of ligand and target proteins

To understand the hydropathy index of the ligand and target proteins, a hydropathy plot was generated using ProtScale (https://web.expasy.org/protscale/). Uniprot IDs of ligand Pyocyanin [Q9HWG9], and two target proteins NOX [P04839] and TLR4 [O00206] were used. Amino acid sequences corresponding to each UniProt ID were retrieved and submitted to the ProtScale server. The Kyte-Doolittle scale and a window scale set to 9 were used for hydropathy plot analysis.

### 5. Molecular Docking

Molecular docking was performed using the HADDOCK (High Ambiguity Driven protein–protein Docking) web server (https://rascar.science.uu.nl/). Ligand and target protein structures retrieved were preprocessed and subsequently renumbered to avoid residue indexing conflicts and formatted according to HADDOCK requirements. The ligand structure (pyocyanin), previously prepared and energy-minimized, was modified for HADDOCK by renaming the residue as “LIG” and assigning unique atom names to ensure compatibility with the docking platform. The most appropriate binding pockets of target proteins were selected after analyzing CastP results. These active residues were specified as docking restraints during submission to guide the interaction between the ligand and the receptor. Both the target protein (TLR4 or NADPH oxidase) and ligand (pyocyanin) structures, along with the defined active site residues, were uploaded to the HADDOCK server. The docking protocol generated multiple conformations, and the resulting complexes were evaluated based on HADDOCK scores and binding energies.

### 6. Visualization and Interaction Analysis

The docking output files for NADPH oxidase and TLR4 were obtained from the HADDOCK server in PDB format. Visualization and analysis of the docked complexes were performed using BIOVIA Discovery Studio Visualizer, which enabled examination of the 3D structure and identification of non-bonded interactions, including hydrogen bonds, hydrophobic contacts, and π-π stacking. The docked complex structures were rendered and exported as high-resolution images in PNG format for representation in the results section.

## Results

### 1. Structural visualization of Pyocyanin ligand and NADPH Oxidase and TLR4 target protein

Figure 1. illustrates the 3D structures of the ligand pyocyanin and two target proteins—NADPH oxidase (NOX) and Toll-like receptor 4 (TLR4)—used for molecular docking. The structure of pyocyanin (**Figure 1A**) reveals its characteristic phenazine ring with planar geometry, composed of fused aromatic rings with nitrogen atoms (blue) and a carbonyl oxygen (red). This configuration supports its redox activity and potential for membrane intercalation.

**Fig. 1.**
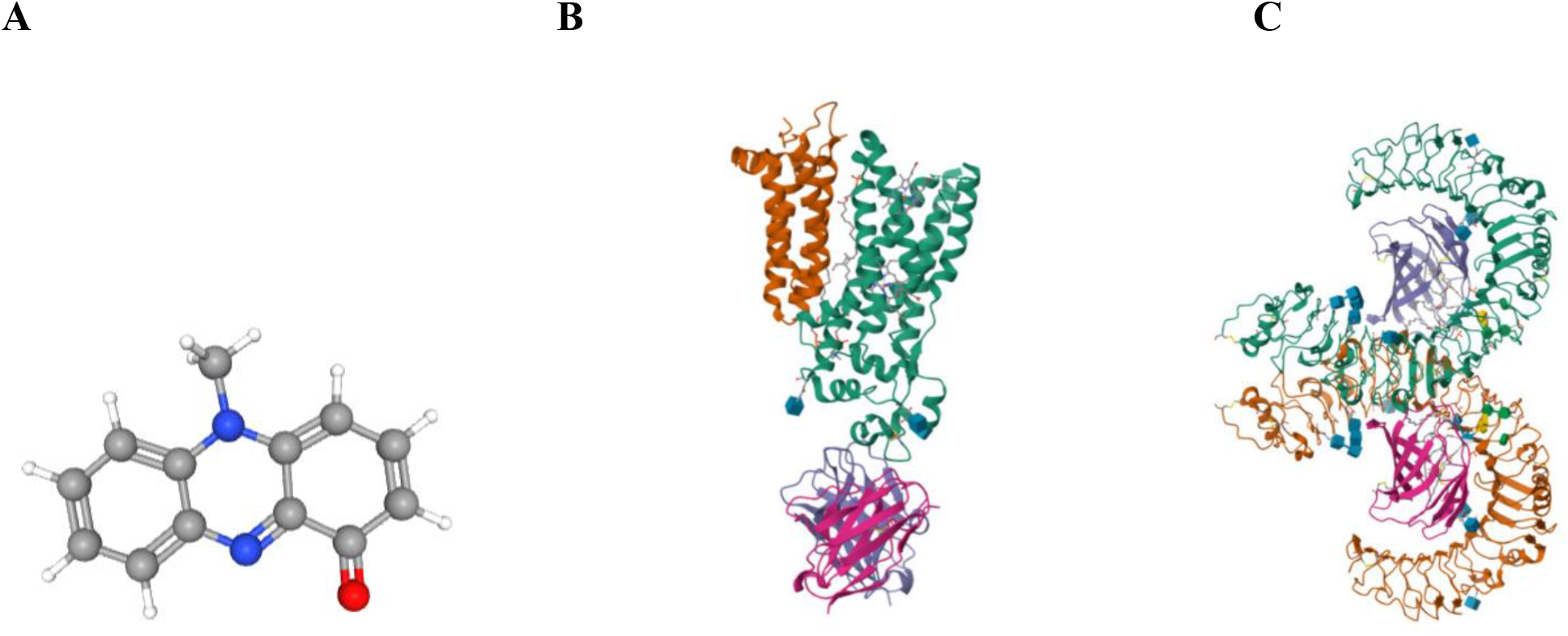
Structure of ligand-Pyocyanin and target proteins-NADPH Oxidase (NOX) and TLR4. **A-** 3D structure of Pyocyanin ligand. Nitrogen atoms are represented by the blue spheres and the red sphere stands for an oxygen atom. **B-** 3D structure of NOX target protein with resolution 3.2 Å (Electron Microscopy). **C-** 3D Structure of TLR4 target protein with 3.1 A^0^ resolution (X-Ray Diffraction). In Both B and C - Alpha-helices are shown as spirals, and beta-sheets are represented by flat arrows.

The NADPH oxidase structure (**Figure 1B**), resolved at 3.2 Å using electron microscopy, shows a complex tertiary organization with distinct alpha-helices and beta-sheets. These structural features are critical for its function in electron transport and ROS generation. Similarly, the TLR4 protein (**Figure 1C)**, resolved by X-ray diffraction at 3.1 Å, exhibits a horseshoe-shaped extracellular domain formed by leucine-rich repeats, important for ligand recognition, along with alpha-helical and beta-sheet secondary structures that stabilize its conformation. The visual representation of both receptor proteins supports their suitability as docking targets due to their defined active sites and well-characterized structural domains.

The binding pocket locations predicted using CASTp are shown in **Supplementary Figures S1.** To identify the most favorable docking sites, the surface topology and potential binding pockets of NOX and TLR4 were analyzed using CASTp (**Figure S1**). The CASTp images reveal clearly defined surface cavities in both proteins, supporting the structural suitability of selected regions for ligand interaction. The binding pockets with the highest volume/surface area was selected for further analysis (**Table S1**).

Hydropathy plots for pyocyanin, NOX, and TLR4 are presented in **Figure S2**. Pyocyanin exhibits an amphipathic profile (Figure **S2A**), with alternating hydrophobic and hydrophilic regions, enabling both membrane interaction and aqueous solubility. The NOX hydropathy profile (**Figure S2B)** reflects its transmembrane nature, with prominent hydrophobic regions stabilizing its membrane anchoring. TLR4 displays variable hydropathy **(Figure S2C**), with alternating peaks and troughs, indicating its potential for both membrane insertion and extracellular ligand interaction.

### 2. Molecular Docking of Pyocyanin with NADPH Oxidase

Molecular docking analysis of Pyocyanin with NOX2 was performed using the HADDOCK platform. Three clusters—Cluster 6, Cluster 3, and Cluster 8—were shortlisted for comparative evaluation based on HADDOCK scores and interaction metrics (**Table 1**).When looking at the docking results for Clusters 6, 3, and 8, Cluster 6 stands out as the most balanced and dependable option. It has the lowest HADDOCK score (84.4 ± 2.9) (**Figure 2A**), which means it shows a stronger bond compared to Cluster 3 (106. 0 ± 3. 5) and Cluster 8 (122. 4 ± 1. 6). The Z-score of –0. 7 also shows that the models in this cluster are among the best in the whole docking run. Cluster 6 also has more similar models, with a cluster size of 13, while Cluster 3 and 8 only have 5 and 4 models respectively, making Cluster 6 more consistent and reliable.

**Table 1:**
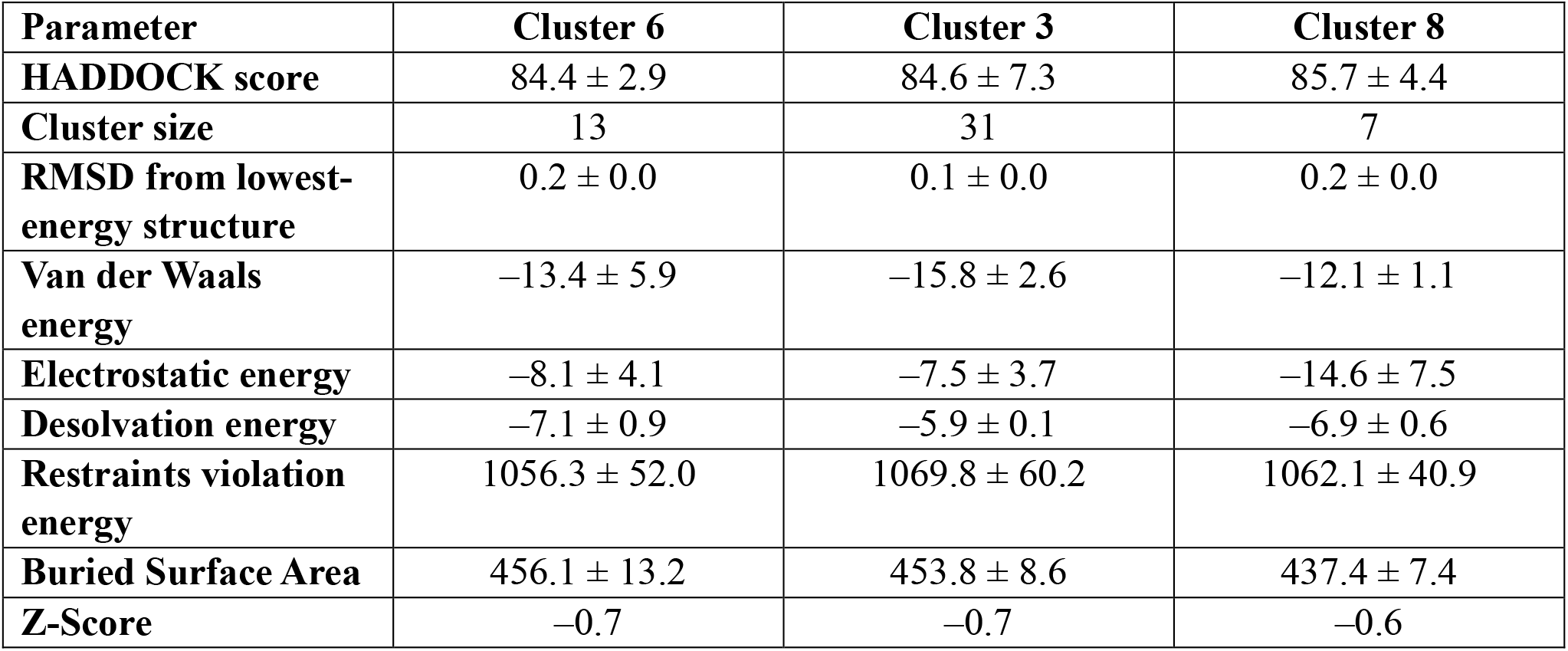
Molecular docking analysis of Pyocyanin with NOX2. Comparison of docking clusters generated by HADDOCK, showing HADDOCK scores, interaction energies, RMSD values, buried surface area (BSA), and Z-scores. Cluster 6 was selected as the most relevant for downstream analysis.

**Fig. 2.**
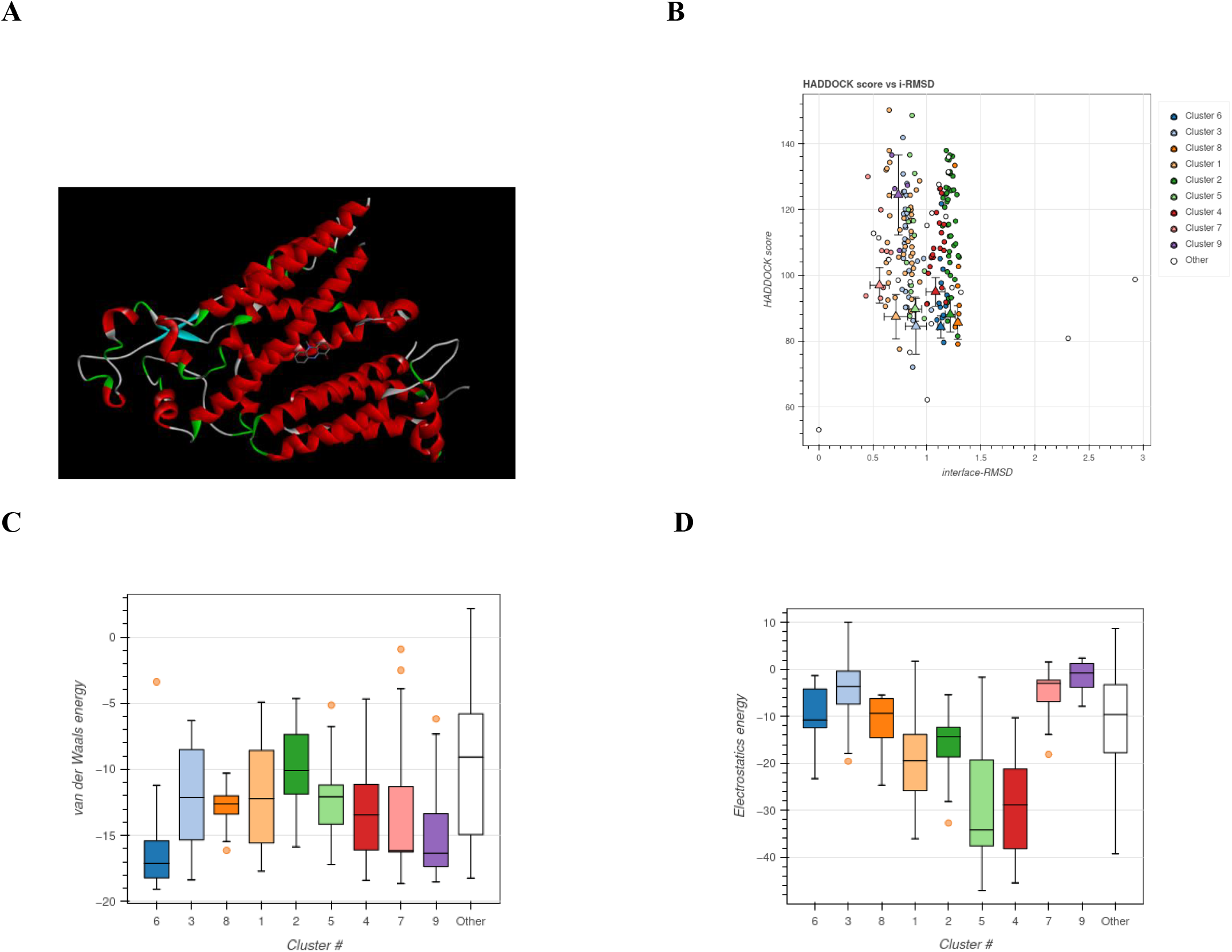
A- Molecular docking pose of pyocyanin with NOX2 (Cluster 6). 3D representation of the docking conformation showing interaction between pyocyanin and the active site residues of NOX2. Pyocyanin is shown bound within the predicted active site of NOX2, with the ligand displayed in stick format. **B- Docking pose distribution for pyocyanin–NOX2 complexes.** Plot of HADDOCK score versus interface RMSD (i-RMSD) depicting the quality and consistency of docking clusters. Cluster 6 shows a tight distribution with low RMSD, indicating pose stability**. C- Van der Waals energy distribution across NOX2–pyocyanin docking clusters**. Box plot showing the spread of Van der Waals energies for each docking cluster. Cluster 3 shows the most favorable VdW interactions. **D- Electrostatic energy distribution across NOX2–pyocyanin docking clusters.** Box plot comparing electrostatic interaction energies across clusters. Cluster 8 displayed the strongest electrostatic interactions.

Among the three clusters analyzed on the HADDOCK platform, cluster 6 shows the lowest HADDOCK score of 84.4 ± 2.9, thereby representing it to be the most stable one among all three clusters. Cluster 6 demonstrated a HADDOCK score of 84.4 ± 2.9, with a RMSD (0.2 ± 0.0) and a buried surface area (BSA) of 456.1 ± 13.2 Å^2^, suggesting a stable and well-defined binding interaction (**Figure 2A and 2B**). The HADDOCK score versus i-RMSD plot (**Figure 2B**) highlights the high pose consistency within Cluster 6, further validating its selection. Despite a high restraint violation energy (>1000), the overall interaction profile of Cluster 6 supports a strong and specific interaction between pyocyanin and NOX2.

Although Cluster 3 had a slightly better Van der Waals energy (–15.8 ± 2.6) and lower RMSD (0.1 ± 0.0), Cluster 6 was chosen for further analysis due to its balanced interaction energies and structural stability. As shown in **Fig. 2C**, for van der Waals interactions, Cluster 6 has the highest value (–13.4 ± 5.9), while Clusters 3 and 8 have weaker ones (–6.7 ± 2.6 and –7.4 ± 1.5, respectively). Cluster 6 shows the most favourable interactions. The same pattern is seen in desolvation energy, where Cluster 6 (–7. 1 ± 0. 9) outperforms Cluster 3 (–1. 9 ± 1. 5) and Cluster 8 (–2. 2 ± 1. 0). Cluster 8, however, has a slight advantage in restraint violation energy, with the lowest value (1003. 0 ± 78. 0), meaning its docking positions fit better with the restraints. But the difference is small, and Cluster 6’s value (1056.3 ± 52.0) is still in an acceptable range. Overall, Cluster 6 offers the best balance.

Electrostatic interaction energy varied across clusters, with Cluster 8 showing the most favorable electrostatics (–14.6 ± 7.5), followed by Cluster 6 (–8.1 ± 4.1) (**Figure 2D**). However, Cluster 6 exhibits the lowest RMSD (0.2 ± 0.0), indicating that its docking positions are tightly packed and stable. Cluster 8 (0.4 ± 0.1) and Cluster 3 (0.5 ± 0.1) show more variation, which could make them less reliable. When it comes to interaction strength, Cluster 6 again performs best, with better electrostatic energy (–8. 1 ± 4. 1) than Cluster 3 (–4. 3 ± 2. 2) and Cluster 8 (–4. 2 ± 1. 8). It’s also worth noting that Cluster 1 (–16. 9 ± 6. 5) and Cluster 2 have even stronger electrostatic interactions, but they don’t perform as well in other areas like docking score and consistency. In short, while Clusters 1 and 2 exhibit strong electrostatic interactions, Cluster 8 has slightly fewer restraint violations and Cluster 3 has a higher HADDOCK score (106.0 ± 3.5), weaker interaction energies, and a smaller cluster size, which makes it less reliable. Cluster 6, however, yields the most well-rounded results. Its combination of strong binding energy, stable structures, and consistent performance across all factors makes it the most reliable choice among the three.

### 3. Molecular Docking of TLR4 with NADPH Oxidase

Molecular docking of pyocyanin with TLR4 was performed using the HADDOCK platform, and the top three docking clusters—Cluster 1, Cluster 4, and Cluster 2—were analyzed for comparative evaluation (**Table 2**).

**Table 2:** Molecular docking analysis of Pyocyanin with TLR4. Comparison of docking clusters generated by HADDOCK, showing HADDOCK scores, interaction energies, RMSD values, buried surface area (BSA), and Z-scores. Cluster 1 was selected as the most relevant for analysis.

| Parameter | Cluster 1 | Cluster 4 | Cluster 2 |
| --- | --- | --- | --- |
| **HADDOCK score** | $25.0 \pm 5.5$ | $30.3 \pm 9.8$ | $33.8 \pm 1.6$ |
| **Cluster size** | 37 | 10 | 18 |
| **RMSD from lowest-energy structure** | $0.3 \pm 0.0$ | $0.3 \pm 0.1$ | $0.4 \pm 0.0$ |
| **Van der Waals energy** | $-15.7 \pm 0.9$ | $-22.7 \pm 1.7$ | $-20.6 \pm 0.6$ |
| **Electrostatic energy** | $-16.9 \pm 6.5$ | $-10.8 \pm 17.6$ | $-7.9 \pm 5.0$ |
| **Desolvation energy** | $-3.0 \pm 0.9$ | $-2.5 \pm 0.4$ | $-3.7 \pm 0.6$ |
| **Restraints violation energy** | $453.5 \pm 54.3$ | $543.5 \pm 124.1$ | $588.6 \pm 15.0$ |
| **Buried Surface Area** | $441.3 \pm 31.4$ | $462.8 \pm 18.4$ | $458.7 \pm 12.1$ |
| **Z-Score** | -2.3 | -1.1 | -0.3 |

Cluster 1 emerged as the most favorable docking solution, with a HADDOCK score of 25.0 ± 5.5, a Z-score of –2.3, and an RMSD of 0.3 ± 0.0, indicating high-quality and statistically significant docking results. The buried surface area (BSA) was 441.3 ± 31.4 Å^2^, suggesting a well-defined and stable binding interface. Additionally, the electrostatic (–16.9 ± 6.5) and van der Waals (–15.7 ± 0.9) interaction energies were highly favorable, supporting strong ligand–receptor interactions (**Figure 3A**).

**Fig. 3.**
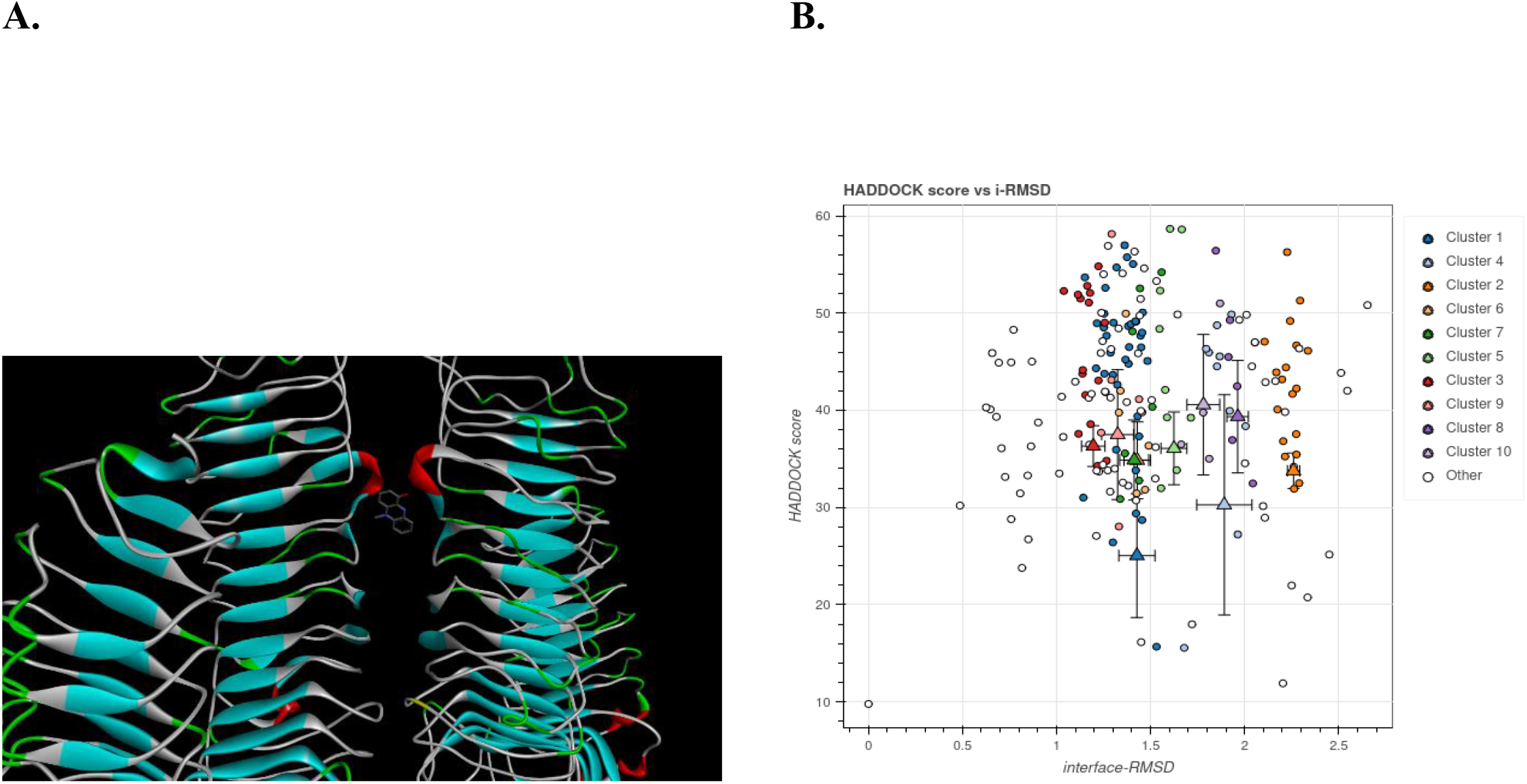
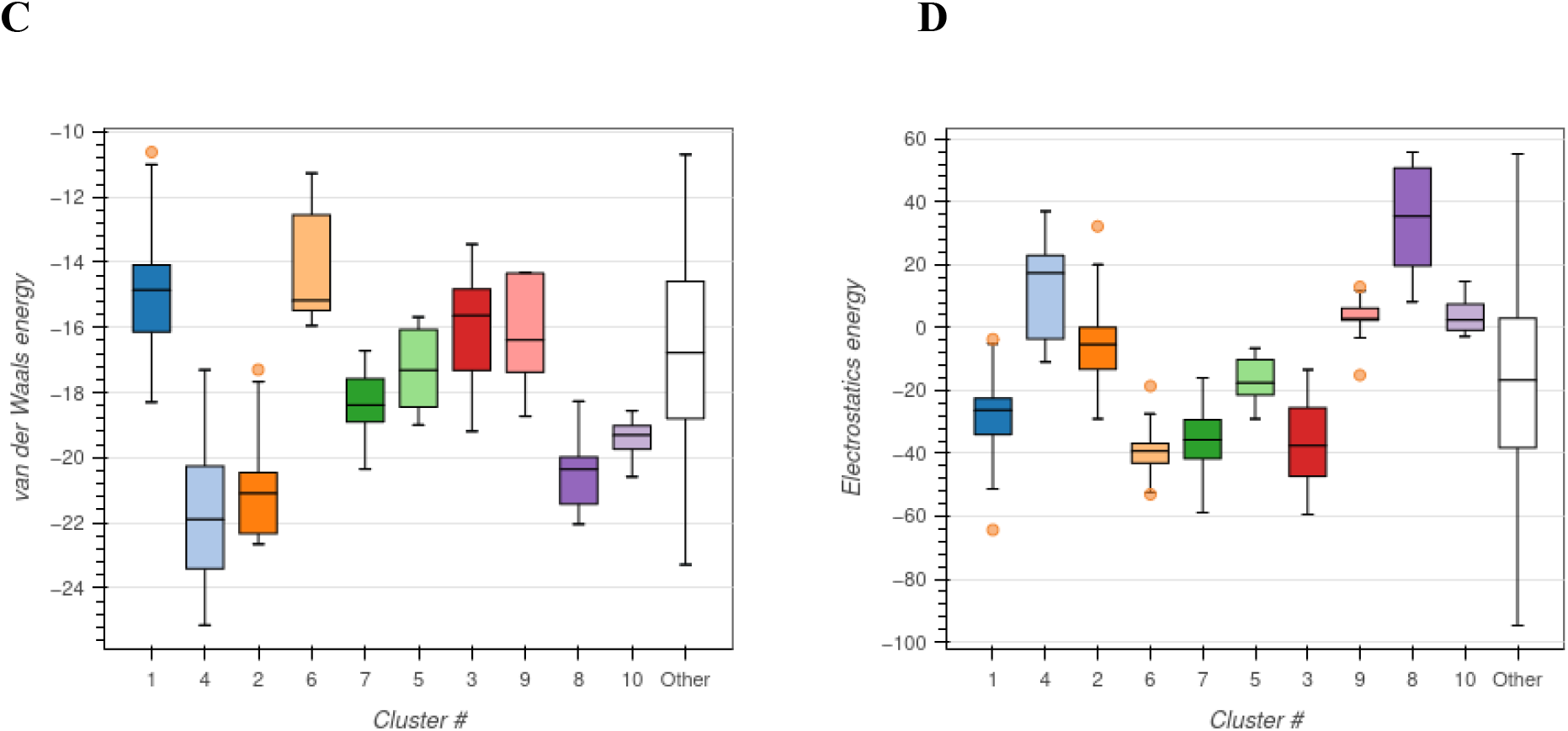
A- Molecular docking pose of pyocyanin with NOX2 (Cluster 6). 3D representation of the docking conformation showing interaction between pyocyanin and the active site residues of NOX2. Pyocyanin is shown bound within the predicted active site of NOX2, with the ligand displayed in stick format. **B- Docking pose distribution for pyocyanin–NOX2 complexes.** Plot of HADDOCK score versus interface RMSD (i-RMSD) depicting the quality and consistency of docking clusters. Cluster 6 shows a tight distribution with low RMSD, indicating pose stability**. C- Van der Waals energy distribution across NOX2–pyocyanin docking clusters**. Box plot showing the spread of Van der Waals energies for each docking cluster. Cluster 3 shows the most favorable VdW interactions. **D- Electrostatic energy distribution across NOX2–pyocyanin docking clusters.** Box plot comparing electrostatic interaction energies across clusters. Cluster 8 displayed the strongest electrostatic interactions.

Although Cluster 4 showed a slightly larger BSA (462.8 ± 18.4 Å^2^) and more favorable van der Waals energy (–22.7 ± 1.7), its higher RMSD (0.3 ± 0.1) and poorer HADDOCK score (30.3 ± 9.8) reduced its overall reliability. Similarly, Cluster 2, with a HADDOCK score of 33.8 ± 1.6, showed weaker electrostatic interactions (–7.9 ± 5.0) and slightly higher restraint violation energy, making it less ideal than Cluster 1.

Visual inspection of the docked complex (**Figure 3A**) revealed that pyocyanin binds within the extracellular domain of TLR4, consistent with its ligand-recognition function. The HADDOCK score vs. i-RMSD distribution plot (**Figure 3B**) confirmed high pose consistency within Cluster 1. Energy distribution plots for van der Waals (**Figure 3C**) and electrostatic interactions (**Figure 3D**) further validated the superior interaction strength of Cluster 1 compared to the other clusters.

Taken together, these findings demonstrate a strong and specific interaction between pyocyanin and TLR4, with Cluster 1 representing the optimal docking configuration for downstream analysis.

### 4. Comparison of molecular docking of Pyocyanin between NOX and TLR4

To assess the relative binding efficiency and interaction characteristics of pyocyanin with the two target proteins, a comparative analysis of the optimal docking clusters—Cluster 6 (NOX2) and Cluster 1 (TLR4)—was performed (**Table 3**).

**Table 3:** Docking Metrics Comparison. Summary of docking results for the most favorable clusters identified by HADDOCK (Cluster 1 for TLR4 and Cluster 6 for NOX2). Shown are HADDOCK scores, Z-scores, cluster sizes, RMSD values, van der Waals energies, electrostatic energies, desolvation energies, restraint violation energies, and buried surface areas (BSA).

| Metric | TLR4–Pyocyanin<br>(Cluster 1) | NOX2–Pyocyanin<br>(Cluster 6) |
| --- | --- | --- |
| HADDOCK Score | 25.0 ± 5.5 | 84.4 ± 2.9 |
| Z-score | -2.3 (excellent) | -0.7 (moderate) |
| Cluster Size | 37 | 13 |
| RMSD (Pose Consistency) | 0.3 ± 0.0 | 0.2 ± 0.0 |
| Van der Waals Energy | -15.7 ± 0.9 | -13.4 ± 5.9 |
| Electrostatic Energy | -16.9 ± 6.5 | -8.1 ± 4.1 |
| Desolvation Energy | -3.0 ± 0.9 | -7.1 ± 0.9 |
| Restraint Violation Energy | 453.5 ± 54.3 | 1056.3 ± 52.0 |
| Buried Surface Area (BSA) | 441.3 ± 31.4 | 456.1 ± 13.2 |

TLR4–pyocyanin interactions exhibited a more favorable HADDOCK score and Z-score, indicating statistically stronger and more reliable docking. TLR4 also demonstrated stronger van der Waals and electrostatic interactions, suggesting more energetically favorable binding.

Conversely, pyocyanin–NOX2 binding exhibited a slightly lower RMSD, indicating higher structural stability in docking poses, and better desolvation energy and BSA, suggesting deeper burial of the ligand in the binding pocket.

Overall, while pyocyanin formed stable complexes with both NOX2 and TLR4, the interaction with TLR4 appears to be stronger and more specific, based on docking metrics. These findings support the hypothesis that pyocyanin may modulate immune signaling pathways via TLR4, while also potentially influencing redox signaling via NOX2.

## Discussion

This study aimed to elucidate the potential molecular interactions between pyocyanin—a virulence factor of *Pseudomonas aeruginosa*—and two host proteins central to oxidative stress and inflammatory signaling: NADPH oxidase (NOX2) and Toll-like receptor 4 (TLR4). Using an in silico molecular docking approach, we demonstrated that pyocyanin can stably interact with both NOX2 and TLR4, suggesting a plausible structural basis for its contribution to oral cancer pathogenesis through ROS generation and immune modulation.

Pyocyanin is a redox-active phenazine that disrupts redox balance by promoting the accumulation of reactive oxygen species (ROS), leading to mitochondrial dysfunction, ER stress, and genomic instability (Kudryavtseva et al., 2016; Reszka et al., 2004). NADPH oxidase is a major enzymatic source of ROS, and its dysregulation has been associated with oxidative damage and chronic inflammation (Tarafdar & Pula, 2018). In our study, molecular docking revealed that pyocyanin interacts stably with the predicted active pocket of NOX2, with Cluster 6 showing the most favorable combination of HADDOCK score, RMSD, and buried surface area. These findings align with previous studies indicating that pyocyanin enhances ROS production via NOX2 by accepting electrons from NADPH (Rada et al., 2013). This interaction may contribute to sustained ROS overproduction, promoting oxidative stress, cellular injury, and tumorigenesis.

Our analysis also showed strong interaction between pyocyanin and TLR4, particularly in Cluster 1, which had a highly favorable Z-score (–2.3) and consistent docking pose. TLR4 is known to be activated by ROS and bacterial components such as lipopolysaccharides (LPS), leading to the initiation of inflammatory signaling pathways through both the TRIF-and MyD88-dependent pathways (González-Carnicero et al., 2023; Ullah et al., 2016). Interestingly, while Cluster 3 of NOX2–pyocyanin and Cluster 8 of TLR4–pyocyanin exhibited more favorable Van der Waals and electrostatic interaction energies, respectively, Cluster 6 (NOX2) and Cluster 1 (TLR4) offered the best overall docking profiles in terms of structural stability and interface quality. This highlights the importance of evaluating multiple docking parameters to select biologically relevant models. ROS produced via NOX2 may amplify TLR4-mediated inflammatory signaling, establishing a positive feedback loop that sustains inflammation and immune dysregulation—hallmarks of the tumor microenvironment in oral cancer.

The involvement of the oral microbiome, particularly *P. aeruginosa*, in carcinogenesis has gained increasing attention in recent years. Pyocyanin’s ability to disrupt immune function and promote a pro-oxidant state may explain how *P. aeruginosa* contributes to oral cancer progression. Our findings provide molecular-level insight into this mechanism, complementing prior in vivo and in vitro studies.

### Limitations and Future Directions

While molecular docking provides valuable insights into ligand–protein interactions, it does not account for protein flexibility, solvent effects, or cellular context. Further validation using molecular dynamics simulations could provide more dynamic insight into the stability of the pyocyanin–NOX2 and pyocyanin–TLR4 complexes. In vitro and in vivo studies are also necessary to confirm the biological relevance of these interactions and to explore their downstream effects on oxidative stress, inflammation, and cancer-related signaling. Additionally, expanding the docking analysis to include mutated forms of NOX2 or TLR4 commonly found in cancer, or other components of the TLR4–NF-κB axis, may offer a more comprehensive understanding of how microbial metabolites shape the tumor microenvironment.

## Conclusion

This study provides novel insights into the molecular interactions between pyocyanin, a redox-active virulence factor of *Pseudomonas aeruginosa*, and two key host proteins—NADPH oxidase (NOX2) and Toll-like receptor 4 (TLR4)—implicated in oxidative stress and inflammatory signaling. Through molecular docking analysis, we demonstrate that pyocyanin binds stably to both NOX2 and TLR4, supporting its potential role in enhancing ROS production and activating immune pathways that contribute to oral cancer pathogenesis. These findings offer a mechanistic framework linking oral microbiome dysbiosis to tumor initiation and progression via host–microbe interactions.

---

**Fig. S1.**
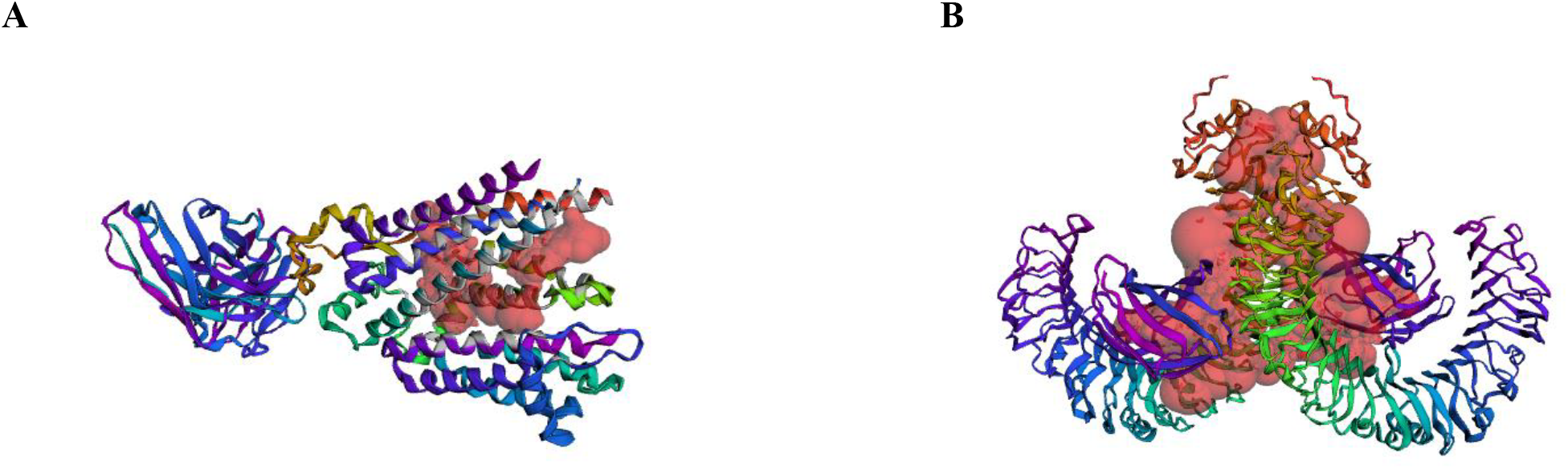
CastP analysis of target proteins. **A-** Surface topology of NOX protein showing predicted pockets. **B-** Surface topology of TLR4 protein showing predicted pockets.

**Fig. S2.**
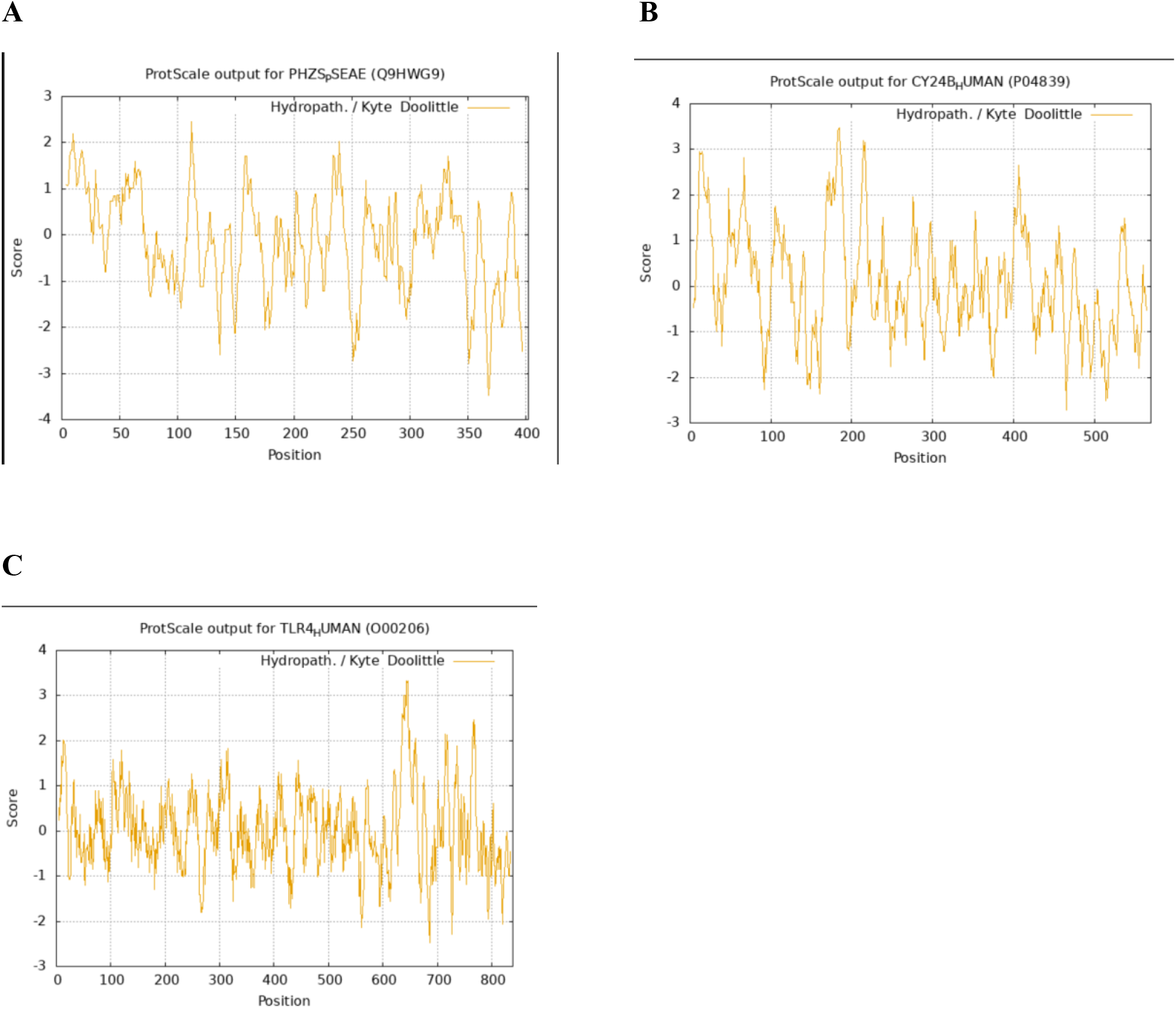
Hydropathy plots of ligand and target proteins. **A-** Hydropathy Plot of Pyocyanin showing alternating hydrophilic and hydrophobic regions, **B-** NOX hydropathy profile reflecting its membrane-bound nature, and **C-** TLR4 hydropathy profile with distinct transmembrane and extracellular regions

**Table S1:** Predicted binding pockets of NOX and TLR4. **A-** Predicted binding sites on NOX. **B-** Predicted binding sites on TLR4

| Target Protein | Area (SA) ($\text{\AA}^3$) | Volume (SA) ($\text{\AA}^3$) |
| --- | --- | --- |
| NOX | 1359.558 | 789.022 |
| TLR4 | 9540.015 | 50536.330 |

